# KMT2C deficiency promotes APOBEC mutagenesis and genomic instability in multiple cancers

**DOI:** 10.1101/2022.02.04.478993

**Authors:** Xiaoju Hu, Antara Biswas, Subhajyoti De

## Abstract

Histone methyltransferase KMT2C is among the frequently mutated epigenetic modifier genes in cancer. It has additional roles in DNA replication, but the effects of KMT2C deficiency on genomic instability during tumorigenesis are unclear. Analyzing 9,663 tumors from 30 cohorts, we report that KMT2C mutant tumors have a significant excess of APOBEC mutational signatures in several cancer types. We show that KMT2C deficiency promotes APOBEC expression and deaminase activity, and compromises DNA replication speed and delays fork restart, facilitating APOBEC mutagenesis targeting ssDNA near stalled forks. APOBEC-mediated mutations primarily accumulate during early replication, and tend to cluster along the genome and also in 3D nuclear contexts. Excessive APOBEC mutational signatures in KMT2C mutant tumors correlate with elevated genomic instability and signatures of homologous recombination deficiency. We propose that in multiple cancer types KMT2C deficiency is a likely driver of APOBEC mutagenesis, which promotes further genomic instability during cancer progression.

## INTRODUCTION

Histone Lysine Methyltransferase 2C (KMT2C), also known as myeloid/lymphoid or mixed-lineage leukemia protein 3 (MLL3) is among the most frequently mutated cancer genes in major cancer types[1–4]. KMT2C is a member of the ASC-2/NCOA6 complex (ASCOM), that is involved in transcriptional coactivation and has histone methylation activity[5–7]. Although type 2 lysine methyltransferases have substantial homology in sequences and closely related functions, KMT2C appears to also have an additional role in DNA replication and genome maintenance[3,8,9]. It has been shown that TP53 recruits MRE11 to sites of stalled replication forks in a KMT2C-dependent manner, and failed restart of replication fork can impair genome maintenance[10]. Furthermore, KMT2C downregulation leads to extensive changes in the activity of DNA damage response and DNA repair genes, and compromises homologous recombination-mediated double-strand break DNA repair, such that treatment with the PARP1/2 inhibitor olaparib leads to synthetic lethality[8]. However, the impact of KMT2C deficiency on mutational landscape and genomic instability at a genome-wide scale in tumor genomes is unclear. Here, using analysis of genomic data from multiple cancer cohorts and biochemical assays, we examine how KMT2C deficiency compromises replication-dependent genome maintenance processes in cancer. We propose that KMT2C mutations may be a key contributor to APOBEC mutagenesis and targetable genomic instability in multiple cancer types.

## RESULTS

### KMT2C mutations frequently occur in multiple cancer types

KMT2C is a relatively large gene with the primary transcript coding a peptide of 4,911 amino acids long and spanning 59 exons. We analyzed mutation status of histone methyltransferases in 7,574 samples from 17 major cancer types from the TCGA[11] and 2,089 samples from 13 ICGC cohorts[12] (Figure 1A; see Methods). Proportion of samples with KMT2C mutations varied between 3.23% (11/364) in liver cancer (TCGA-LIHC) and 40% in colorectal cancer (ICGC-COCA-CN) and lung squamous cell carcinoma (ICGC-LUSC-KR). A majority of these mutations are missense, nonsense and frameshift mutations (Figure 1B) with potentially loss-of-function consequences, as reported elsewhere[5,8]. While the mutations are distributed throughout the gene locus, there are two mutational hotspots around exon 36-38, and exon 43-52, closer to the 3’ end of KMT2C gene in multiple cancer types (Figure 1B). Exons 36-38 and 43-52 both overlap with the extended PHD zinc finger domain annotation, while 3’ side of exon 52 also overlaps with the FY-rich domain.

**Figure 1.**
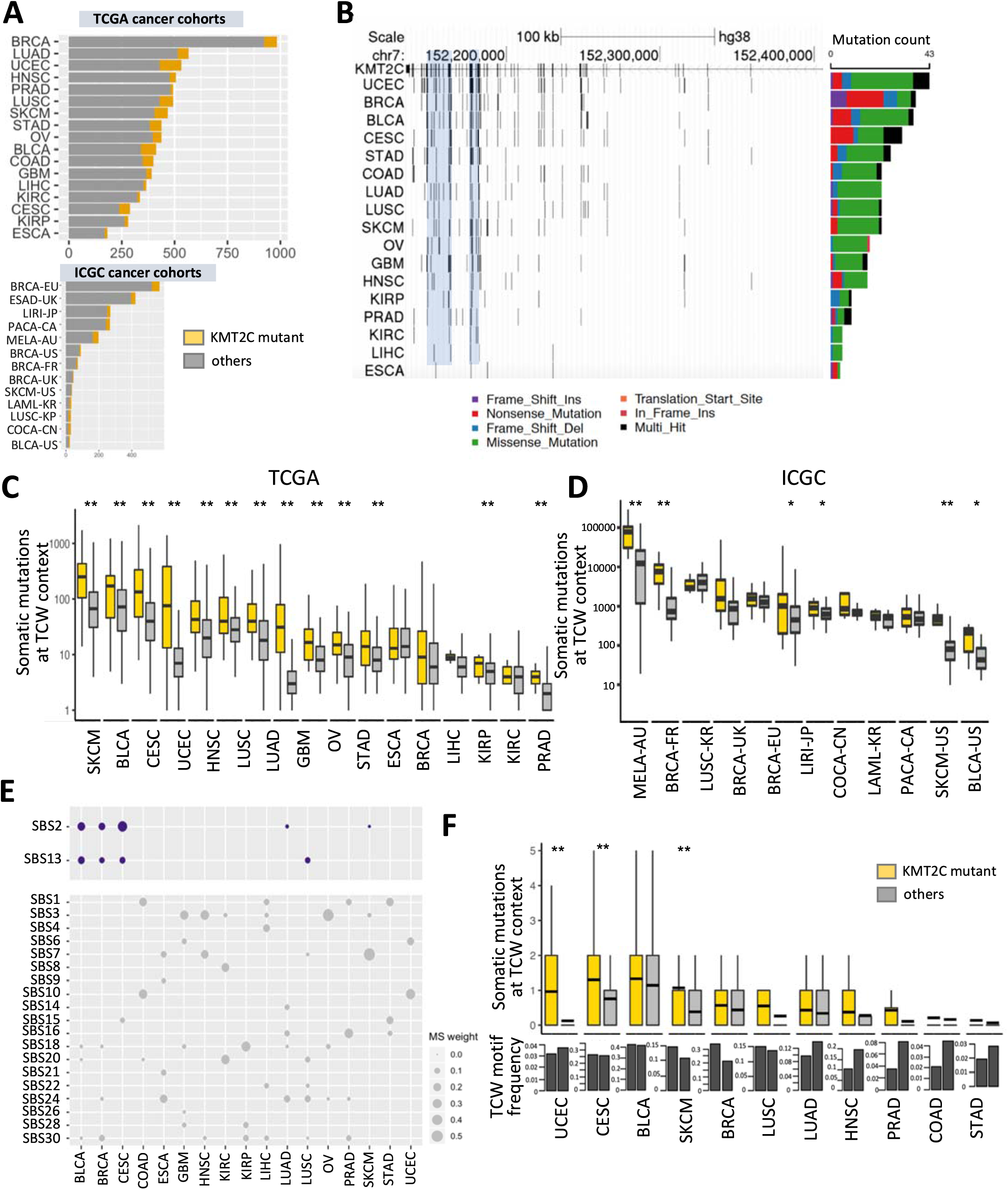
KMT2C mutant tumors have a significant excess of APOBEC mutational signatures in multiple cancer types. (A) The number of tumor samples in 17 TCGA and 13 ICGC cancer cohorts. The samples with KMT2C mutations are indicated in yellow. (B) Genome graph shows distribution of non-silent, coding mutations in KMT2C locus in the TCGA cancer cohorts. The shadowed blue regions indicate two mutational hotspots spanning exons 36-38 and 43-52. The bar graph on the right depicts total frequency of different classes of mutations in respective cohorts, as indicated by the keys. (C-D) Boxplot showing the somatic mutation count at TCW contexts (including TCA to TTA or TGA, and TCT to TTT or TGT mutations) in KMT2C mutant (yellow) and other samples (grey) for (C) TCGA and (D) ICGC cohorts. P-value was calculated by Mann–Whitney U-test, *p-value<0.05, **p-value < 0.01. (E) Bubble plot showing the weights of different COSMIC v2 mutational signatures in non-silent KMT2C mutations in respective cancer cohorts. (F) Boxplot showing the somatic mutation count at TCW contexts within ATAC-seq peak regions in KMT2C mutant and other samples in the TCGA cohorts. Barplots in the panel below show the average frequency of TCW motifs in these ATACseq regions in the two groups in these cancer cohorts. P-value was calculated by Mann–Whitney U-test, *p-value < 0.05, **p-value < 0.01. For description of the cohorts, see Figure 1A.

### KMT2C mutant tumors have an excess of APOBEC mutational signatures

Given the dual role of KMT2C in both epigenetic regulation and replication, we first examined whether somatic mutations in KMT2C deficient tumors have an excess of signatures of certain mutational processes that could in turn provide mechanistic insights into the effects of loss of KMT2C on genome maintenance. In each cancer cohort, we identified the samples with non-silent mutations in KMT2C (see Methods) and designated them as KMT2C-mutant tumors. We then examined the single base substitution patterns characteristic of different mutational processes in somatic mutations in the tumor samples in the cancer cohorts.

APOBEC is a common source of somatic mutations that arise during replication stress in cancer genomes and preferentially occurs at TCW trinucleotide context (W = A or T; specifically, TCA to TTA or TGA, TCT to TTT or TGT)[13]. Comparing frequencies of these somatic mutation-classes between the KMT2C mutant and other samples in different TCGA[11,12] and ICGC cancer cohorts, we observed that the former had a significant excess of APOBEC-mediated mutations in a majority of the cohorts (Figures 1C and 1D). This trend was especially prominent in the cohorts with high baseline APOBEC mutational signatures such as bladder cancer (TCGA-BLCA and ICGC-BLCA-US), cervical cancer (TCGA-CESC), head and neck cancer (TCGA-HNSC), lung cancer (TCGA-LUSC and TCGA-LUAD), and breast cancer (ICGC-BRCA-FR and ICGC-BRCA-EU). Next, using an unbiased approach we examined all COSMIC (v2) mutational signatures in all the samples from the TCGA and ICGC projects. Among the mutational signatures prominently present (>5% mutational weight) in both datasets, APOBEC mutational signatures SBS2 and SBS13 showed systematic increase in mutational weight in KMT2C mutant samples (Figure S1A). Subsequently, in a cohort-wise analysis we calculated the proportional weights of each mutational signature in each cancer cohort, and observed that APOBEC mutational signature (combined mutational weight of SBS2 and SBS13) was substantially higher in KMT2C mutant tumors, especially in the cohorts that had reasonable footprints of APOBEC mutagenesis, including cervical cancer (TCGA-CESC), bladder cancer (TCGA-BLCA), head and neck cancer (TCGA-HNSC), lung cancer (TCGA-LUSC) and breast cancer (ICGC-BRCA-EU) (Figures S1B and S1C). Taken together, enrichment for APOBEC mutational signature was consistently observed in both the TCW context- and mutational signature-guided analyses. Besides the APOBEC mutational signatures, several other mutational signatures were also detected in the pan-cancer and cohort-wise analyses, but their effects were moderate and/or cohort-specific; in the pan-cancer analysis SBS6, SBS7, SBS10 and SBS12 showed over-representation in KMT2C mutant tumors (Figure S1A), while in the cohort-wise analysis SBS20, which is attributed to defective DNA mismatch repair (MMR) was also enriched in the KMT2C mutant samples (Figure S1C). KMT2C deficiency contributes to replication stress[14], which promote microsatellite instability (MSI) and hypermutation.

APOBEC family enzymes APOBEC3A (A3A) and APOBEC3B (A3B) are potent mutator deaminases, which contribute to the characteristic APOBEC mutational signatures. But A3A and A3B have differences in the tetranucleotide context preferences for deamination: APOBEC3A prefers YTCA, while APOBEC3B prefers RTCA (Y = pyrimidine, R = purine) motifs[15]. Furthermore, APOBEC3B has much higher expression level than APOBEC3A[16]. In our analysis, APOBEC-mediated mutations at both the RTCA and YTCA contexts showed increased incidence in KMT2C mutant tumors, suggesting that the associations are probably not specific to APOBEC3A or APOBEC3B-mediated mutagenesis (Figure S1D).

To examine whether APOBEC-mediated or other mutagenic processes led to KMT2C mutations in the cancer genomes, we determined the burden of different mutational signatures on the non-silent mutations in KMT2C in each cohort. There were no single, universal mutagenic processes driving KMT2C mutations across different cancer types, albeit SBS1, SBS3, SBS7, and SBS10 were observed in multiple cancer types. SBS2 and SBS13 - the signatures of APOBEC mutagenesis were observed in some TCGA cancer cohorts (e.g., BLCA, BRCA, CESC, SKCA, LUSC; Figure 1E), which had high prevalence of APOBEC-mediated mutations. In some cases, these mutations are causal events, while in other once a loss-of-function mutation is acquired, the non-functional allelic copy might continue to accumulate additional mutations under the prevailing mutagenic processes in the affected samples, but we could not ascertain the order of mutational events.

### Patterns of mutational signatures in epigenomic contexts

Given the role of KMT2C as an epigenetic modifier, we first assessed whether APOBEC mutagenesis predominantly occurs within the altered chromatin domains in the KMT2C deficient tumors. We jointly analyzed somatic mutations and ATAC-seq data from the same samples from the TCGA cancer cohorts, and observed that KMT2C mutant tumors had higher burden of APOBEC mediated mutations in ATAC-seq peak regions, when compared to that in the ATAC-seq peak regions in other samples in the same cohorts. This was also applicable to a majority of the cohorts with high prevalence of APOBEC-mediated mutations such as breast cancer (TCGA-BRCA), bladder cancer (TCGA-BLCA), cervical cancer (TCGA-CESC), lung squamous cell carcinoma (TCGA-LUSC), which also had high frequency of TCW motifs in the ATACseq peak regions (Figure 1F). In a complementary analysis, we overlaid open chromatin data from KMT2C null and proficient HTB9 bladder and MCF7 breast cancer cell lines with somatic mutations from TCGA tumors, and found that altered chromatin regions gained in KMT2C null conditions had excess of somatic mutations at TCW trinucleotide context in multiple cancer types (Figure S2A). However, a vast majority of the APOBEC mediated mutations occurred outside the peak regions (Figure S2A), and in both analyses the increased burden of APOBEC-mediated mutations was not restricted only to the altered chromatin domains but observed genome-wide - led us to investigate mutagenesis in different genomic, epigenomic, nuclear, and replication contexts systematically.

Therefore, next we broadened the analysis, segmenting the genome into a number of contexts that are relevant for key cellular processes such as transcription, regulation of gene expression and DNA replication, and calculating enrichment for the mutational signatures in these contexts in the KMT2C mutant tumors (Figures 2A and S2B). The repeat and gene regions showed enrichment for APOBEC-mediated mutations (SBS2 and SBS13), but within gene regions, exons did not show such preferences. There were other, signature-specific differences as well. For instance, SBS13, which is known to occur more often in early replicating[17], open chromatin context showed higher burden in euchromatin context in KMT2C mutant tumors in BRCA-EU cohort. Interestingly, SBS2, which preferentially occurs in heterochromatic, late replication contexts, also showed relative enrichment in all chromatin contexts in KMT2C mutant tumors in this cancer type (Figure 2A). Some other differences were cancer-type specific (Figure S2B). Taken together, in KMT2C mutant tumors, APOBEC mediated mutations occurred not just in altered chromatin domains but genome-wide and showed preferences for broader [18] genomic contexts - motivating us to investigate the observed association in the context of the role of KMT2C in DNA replication.

**Figure 2.**
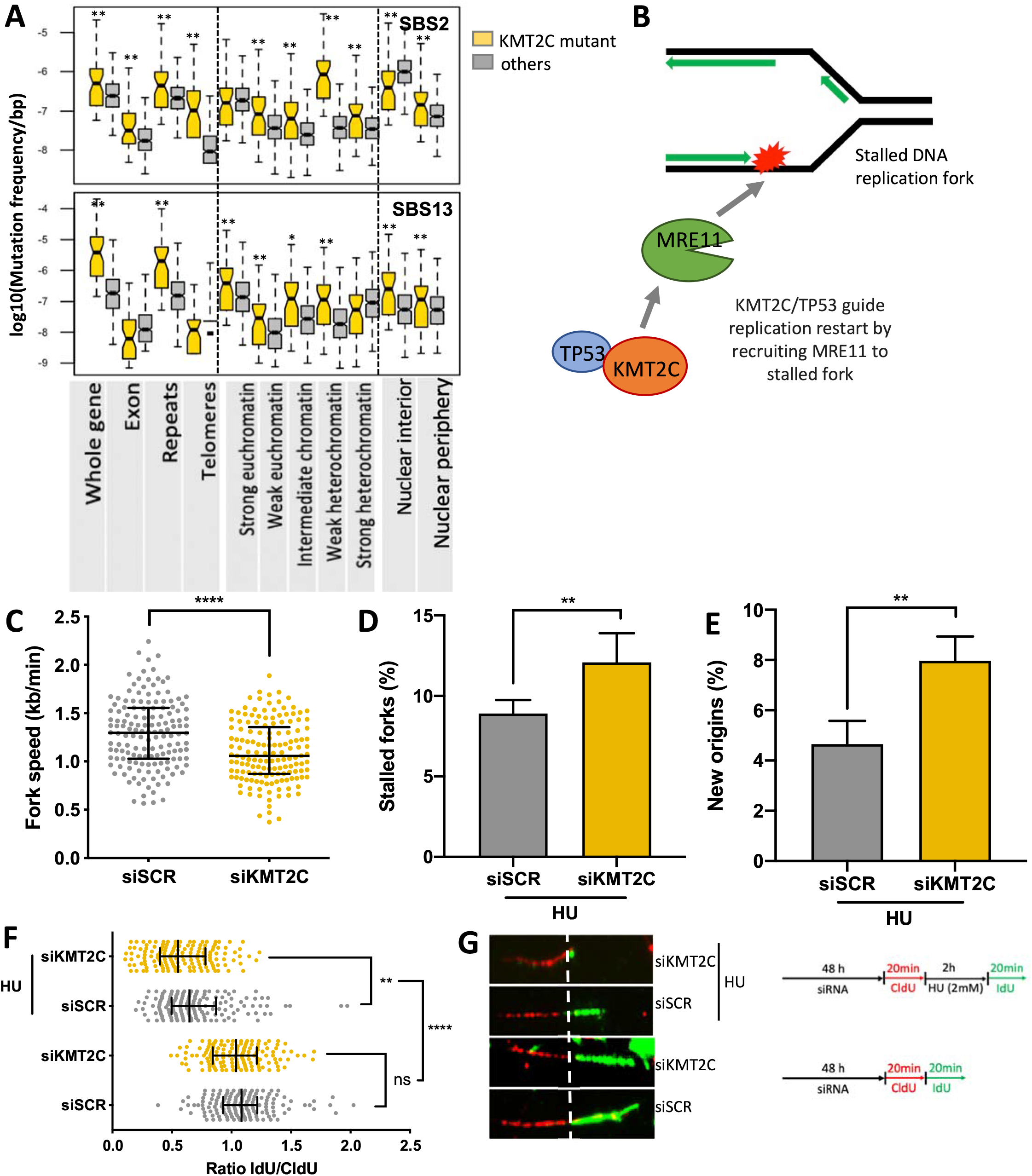
KMT2C deficiency is associated with defects in replication. (A) Mutational burden of SBS2 and SBS13 mutational signatures in different genomic, epigenomic, and nuclear localization contexts in KMT2C mutant (yellow) and other (grey) samples in the breast cancer cohort (BRCA-EU). Similar results for other cohorts are shown in Figure S2A. P-value was calculated by Mann–Whitney U-test, *p-value < 0.05, **p-value < 0.01. (B) Schematic representation showing the paradigm that KMT2C/TP53 guide replication restart by recruiting MRE11 to stalled DNA replication fork. (C-G) DNA fiber assay on control (siSCR) and KMT2C knockdown (siKMT2C) HEK293T cells showing fork speed with line and bars representing the median and interquartile range (C), % of stalled forks with error bars representing SEM (D), % of newly initiated origins with error bars representing SEM (E), ratio of IdU/CldU with line and bars representing the median and interquartile range (F) and representative images of DNA fibers with a schematic representation of the DNA fiber assay conditions (G). Experiments performed with or without hydroxyurea (HU) treatment are indicated in the schematic. P-value was calculated using Mann–Whitney U-test wherein *p-value < 0.05, **p-value < 0.01, ***p-value < 0.001, ****p-value < 0.0001.

### KMT2C loss causes replication stress

KMT2C plays a key role in recruitment of MRE11 to restart stalled DNA replication fork [9] facilitating replication and genome maintenance (Figure 2B). Stalled replication forks and unscheduled origin firings produce ssDNA, which can be potential targets of APOBEC mutagenesis. Thus, we examined whether KMT2C deficiency affects DNA replication dynamics. We first evaluated replication fork dynamics in HEK293T cells using DNA fiber assay, which allows visualization and analysis of replication at DNA-strands level. We knocked down the expression level of KMT2C (siKMT2C) in these cell lines, labelled them with CldU (red tracks) followed by IdU (green tracks) and measured frequency and lengths of fiber tracks. Cells transfected with Scrambled siRNA (siSCR) served as a control. Under unperturbed growth conditions, KMT2C deficiency caused DNA replication stress in HEK293T cells, as indicated by shorter nascent tract length (Figure S2C) and decreased replication fork speed (Figure 2C). Earlier studies have reported implication of KMT2C-mediated chromatin opening in MRE11 nuclease recruitment to stalled forks. We reasoned that apart from contribution to downstream replication fork maintenance KMT2C also functions upstream to prevent DNA replication stress. To further dissect the role of KMT2C in DNA replication, we treated cells with hydroxyurea, a genotoxic agent which depletes nucleotide pools, as a prototype of replication stress. We prelabeled cells with CldU prior to HU treatment, followed by IdU and determined the length and frequency of ongoing replication forks stalled upon hydroxyurea treatment, as well as licencing of new/dormant origins, if any. We observed an increase in frequency of stalled forks in KMT2C silenced cells challenged by HU (Figure 2D). Increased fork stalling in these cells was compensated by an increase in aberrant origin firing, which may impede progression of already established forks (Figure 2E). For the stalled forks that could resume DNA synthesis, KMT2C knocked down cells exhibited defective replication resumption compared to wild-type cells as assessed by the ratio of IdU to CldU tract lengths (Figures 2F, 2G). These observations are consistent with reports that recruitment of MRE11 on stalled replication forks to initiate DNA repair is dependent upon KMT2C mediated chromatin opening [9,19], and suggest that KMT2C deficiency invokes replication stress and disables the cells for efficient replication restart and fork progression upon genotoxic stress. We further observed MRE11 and KMT2C expression significantly associated with mutational signatures of APOBEC mutagenesis (DBS11) [18] and replication stress (SBS40) [20]in the PCAWG pan cancer dataset (Figure S3A), which led us to examine APOBEC expression and activity in KMT2C deficient conditions.

### Replication stress triggers APOBEC activity

Among APOBEC family genes, APOBEC3A or APOBEC3G have almost undetectable base-line expression whereas APOBEC3B remained detectable in most human cell lines[15]. We performed RT–qPCR analysis in KMT2C knockdown HEK293T, RPE1 and MDA-MB-231 cells with low, moderate and high APOBEC3B expression levels, respectively (Figure S3B). Cells transfected with non-targeting scrambled siRNA (siSCR) served as a control. Figure 3A shows the levels of KMT2C, MRE11 and APOBEC3B in HEK293T cells, with small but insignificant reduction in MRE11 and unchanged APOBEC3B expression levels in KMT2C deficient cells as compared to control cells. However, when KMT2C depleted cells were challenged with DNA replication inhibitor HU, significant reduction in MRE11 expression and induction in APOBEC expression compared to that in the control cells were observed. Surprisingly, in RPE1 and MDA-MB-231 cells, KMT2C silencing resulted in MRE11 depletion and APOBEC induction, without additional replication stress. Overall, we observed a trend of elevating APOBEC3B levels upon KMT2C loss in all three cell lines, HEK293T, RPE1 and MDA-MB-231 cells with varying levels of APOBEC3B expression. Integrating transcription factor binding sites and ATAC-seq data for the TCGA cancer cohorts we observed that (i) KMT2C mutant tumors have relatively more open chromatin near well-characterized regulatory regions [21] around APOBEC3B promoter, (ii) these regulatory regions overlap with the binding sites of the NF-κB and AP-1 family of transcription factors, which are known regulators of APOBEC3B, and (iii) such chromatin-level alterations are also associated with increased APOBEC3B mRNA expression in the same samples, as observed in multiple cancer cohorts (Figure S3D).

**Figure 3.**
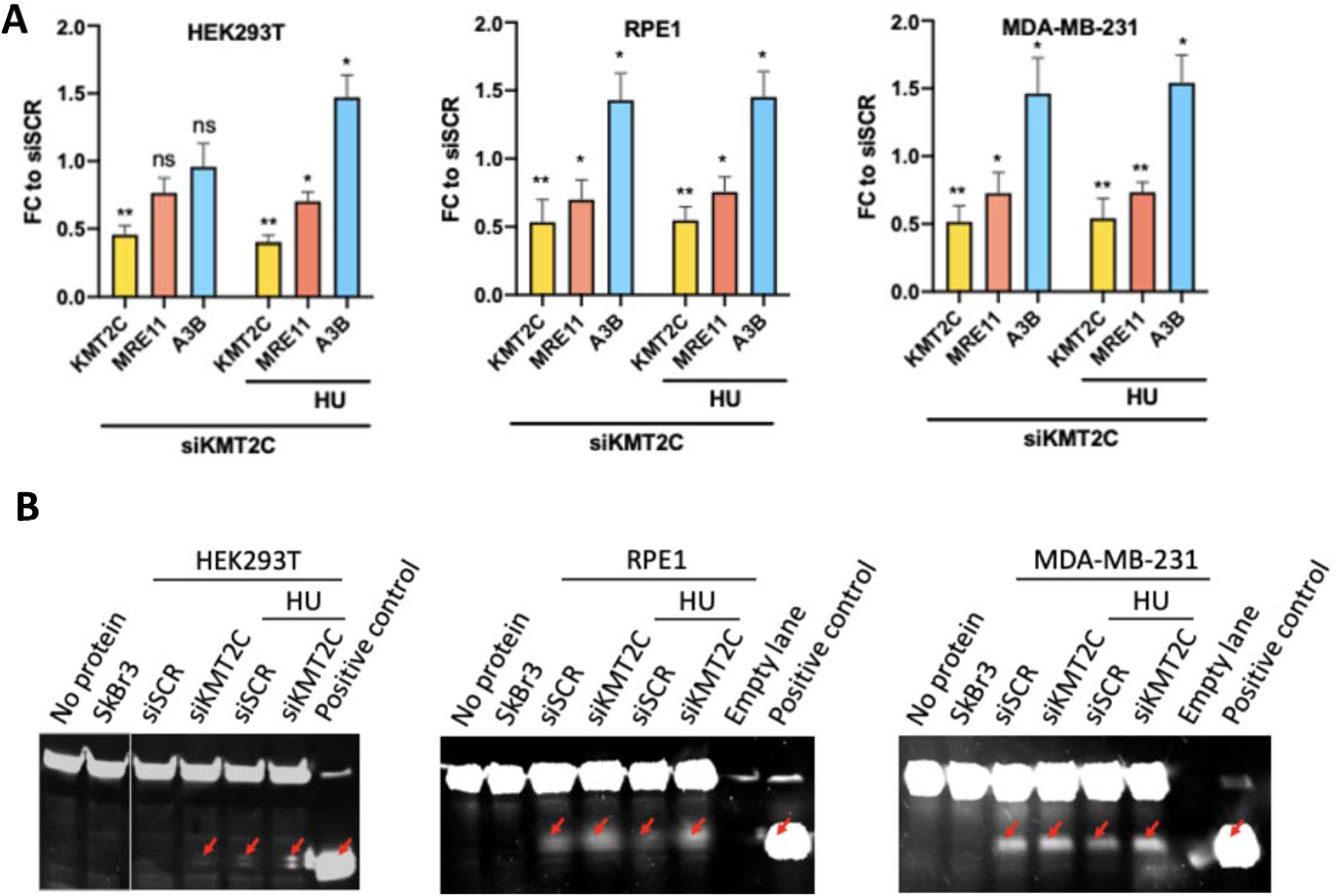
Replication stress due to KMT2C deficiency promotes APOBEC expression and deaminase activity. (A) Fold change (FC) in expression of indicated genes in HEK293T, RPE1 and MDA-MB-231 cells in response to KMT2C loss (siKMT2C) and/or HU treatment as compared to control cells (siSCR) with error bars representing SEM. Asterisks indicate statistical significance. P-value was calculated using t-test wherein *p-value < 0.05, **p-value < 0.01. (B) Protein lysates were assessed in deaminase assay. Test probe without protein lysate and lysate from SkBr3 cell line were used as negative controls. Probe containing uracil rather than cytosine and not incubated protein lysate, showing the mobility of the cleaved product was used as positive control (last lane). Red arrows indicate cleaved probe.

Although the base-line expression of APOBEC3B in human cells is generally low, and elevated expression is associated with increased APOBEC activity[22], we further examined whether KMT2C loss triggers elevated levels of APOBEC deaminase activity as well. We examined the deamination activity present in KMT2C deficient cell lysates using an oligonucleotide-based cytidine deamination assay where C to U conversion in labeled oligonucleotide allows fluorescence detection following cleavage by uracil-DNA glycosylase activity. In lysate from SkBr3 cell line, no deaminase activity was detected, an APOBEC3B-null control for this assay (Figure 3B). Scrambled siRNA transfected HEK293T cells showed no detectable deaminase activity, owing to low baseline APOBEC expression, but in corresponding KMT2C deficient cells deaminase activity was elevated as evidenced by cleaved probe, deaminated by increased APOBEC3 expression in cell extracts, which further increased several folds when subjected to HU treatment. Similar to RT-qPCR results, in RPE1 and MDA-MB-231 cells, upon KMT2C depletion APOBEC3 deaminase activity was elevated, even without genotoxic insults. Quantified deamination results indicating the percentage of the probe cleaved with cell lysate under indicated conditions is shown in bar graphs (Figure S3C). These evidences are collectively consistent with a model that KMT2C loss and replication stress increase APOBEC3 expression and activity, contributing to APOBEC mutagenesis.

### APOBEC mutational signatures in replication context

APOBEC mutagenesis during replication results in mutational signatures SBS2 and SBS13[17], which preferentially occur in late and early replication contexts, respectively (Figure 4A). Analyzing somatic mutations from the breast cancer cohort (BRCA-EU) with repliseq data from MCF7 cell line, we observed that in the tumors with no KMT2C mutations indeed SBS2 mutation density increased from early to late replication, while SBS13 had comparable mutation density through all replication timing domains. In contrast, KMT2C mutant tumors had an excess of both SBS2 and SBS13 mutations in early replication contexts, although the proportional increase was higher for SBS2 (Figure 4B). APOBEC-mediated mutations tend to occur sporadically and in bursts, and in case of replication stress there are clusters of APOBEC-associated mutations which have extended processivity i.e. groups of similar substitutions attributed to the same mutational signature on the same replication strand[16,17]. We calculated processivity of the APOBEC mutational signatures SBS2 and SBS13, and compared that between KMT2C proficient and deficient tumors. The processivity groups where consecutive mutations are within 10kb were considered, since more distant ones may not necessarily occur in the same replication stress-related event. SBS13 was generally associated longer processivity group sizes than SBS2, which is consistent with previous studies[17]. Compared to other tumors, KMT2C mutant tumors had higher frequencies across nearly all processivity groups for SBS2 and SBS13, and had preference for longer processive group events (e.g. n>6), suggesting preference for clustered APOBEC mutagenesis (Figure 4C). Clustered mutagenesis can be associated with kataegis, but we found no significant enrichment for such events in KMT2C mutant tumors.

**Figure 4.**
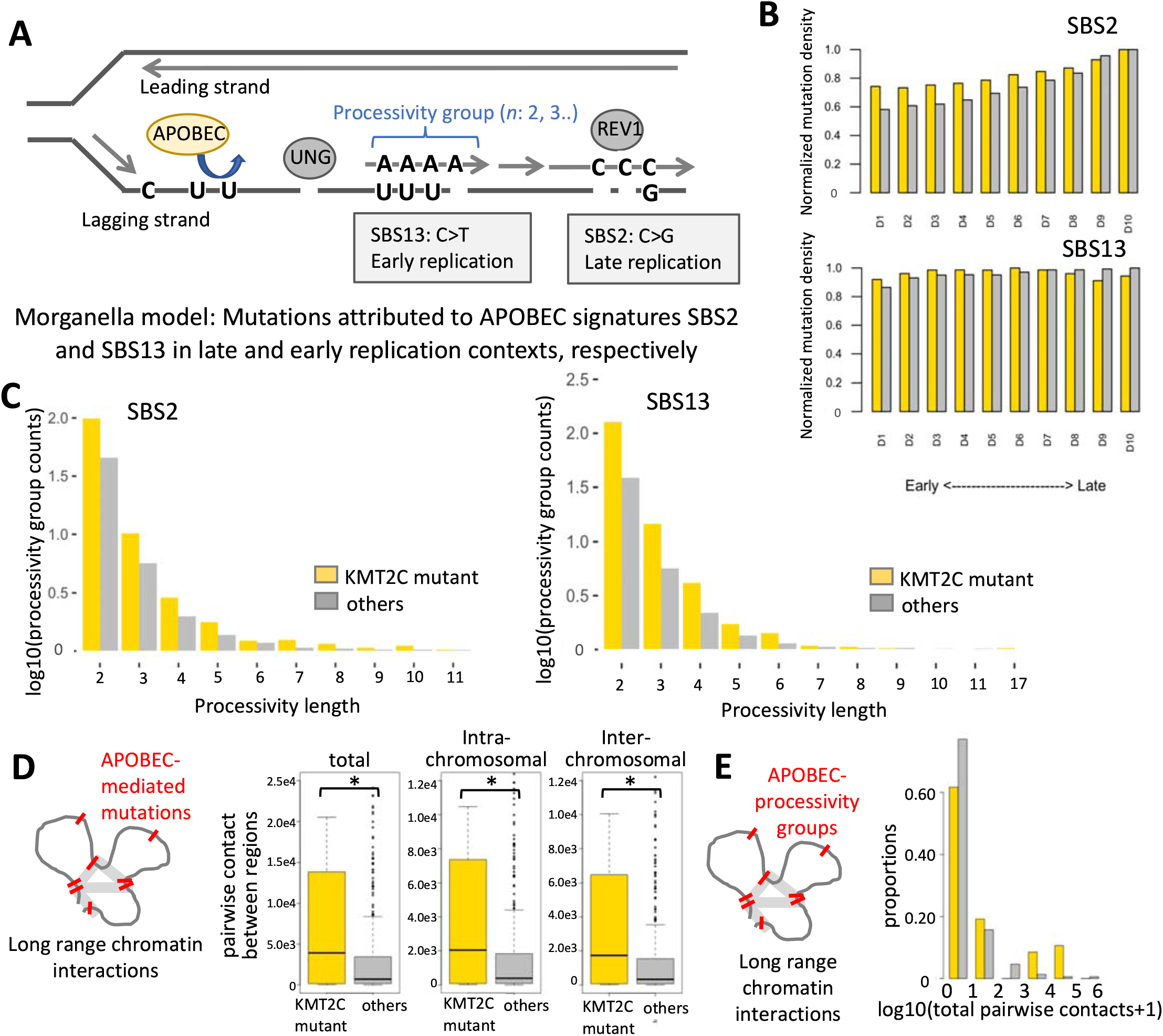
APOBEC mutational signatures in replication contexts. (A) Schematic representation showing that mutations attributed to APOBEC signatures SBS2 and SBS13 preferentially occur in late and early replication contexts, respectively. Such mutations may occur in clusters on the same strand, as indicated by the length of processivity groups. (B) Boxplots showing normalized mutation densities of APOBEC mutational signatures SBS2 and SBS13 in different DNA replication contexts in KMT2C mutant (yellow) and other (grey) samples in the breast cancer cohort (see Methods for details). The frequencies of acquired mutations in respective replication contexts were normalized by the total length of such regions (‘A’s,’C’s,’G’s,’T’s, and excluded ‘N’s). (C) Frequencies of APOBEC processivity group events corresponding to different processive group lengths in KMT2C mutant (yellow) and other (grey) samples in the breast cancer (BRCA-EU) cohort. The frequencies of acquired mutations in each group were normalized by the number of samples in corresponding groups, and nominalized counts were log10 transformed. (D) Boxplots showing the total, intra- and inter-chromosomal pairwise contacts between genomic regions with APOBEC-mediated mutations in KMT2C mutant and other tumors in the BRCA-EU cohort. *p-value < 0.05. (E) Histogram showing the proportions of tumors with different frequencies of pairwise contacts between genomic regions with APOBEC-mediated mutation processive groups in KMT2C mutant and other tumors in the BRCA-EU cohort. The difference was statistically significant (p <0.05).

### Clusters of APOBEC mutagenesis localized in 3D nuclear contexts

Since replication of multiple DNA segments concurrently progress within 3D nuclear organizations called replication factories[23], we examined whether there is APOBEC-mediated mutations cluster within 3D nuclear contexts, even when those might be distal on the linear DNA, especially in the KMT2C mutant tumors. We overlaid HiC-based long-range chromatin interaction data [24] with somatic mutations attributed to APOBEC mutagenesis for the breast cancer (Methods) and observed that KMT2C mutant tumors had significantly higher number of pairwise long-range interactions between genomic regions harboring APOBEC-associated mutations compared to other samples in the cohort, and the overall trends were similar for both intra- and inter-chromosomal interactions (Figure 4D; total, p-value=1.35e-2; intra-chromosomal, p-value=1.40e-2; inter-chromosomal, p-value=1.18e-2). We then repeated the analyses only for a subset of the genomic regions that harbor the APOBEC processivity groups (n≥2) as above, and found that 3D clusters of mutations, marked by excessive number of between-region contacts was significantly more common in KMT2C mutant tumors (Figure 4E; p-value=9.5e-06). This was not due to long processivity groups within the same DNA segments; long-range interaction had a resolution of 100kb, each pair of regions was counted only once per sample, and we observed similar results for within and between-chromosome interactions. Our results suggest that KMT2C mutant tumors accumulate frequent clusters of APOBEC within 3D nuclear contexts. It is plausible that many such mutation clusters may arise concurrently during replication within the same replication factory, but it is challenging to ascertain that conclusively.

### Replication stress skews DNA repair away from canonical HR pathway

Replication stress and delay in fork restart in KMT2C deficient conditions can promote DNA double strand breaks[25,26], and it is suspected that KMT2C loss contributes to defects in homologous recombination, causing sensitivity to PARP inhibitors in cell line models[8]. We quantified the extent of genomic DNA damage by immunostaining KMT2C depleted HEK293T and RPE1 cells with 53BP1 which identifies DNA breaks. We observed that DNA damage repair in KMT2C deficient cells, as evidenced by the number of 53BP1 foci positive cells, number of cells with more than five foci and number of foci per cell, was higher as compared to those measured in the control cells (Figures 5A, 5B and S4A). It is known that 53BP1 does not solely promote NHEJ, but is also required for HR[27,28]. Since the choice of repair pathway is crucial to maintain genome integrity, we asked whether HR or other non-HR pathways might be used to repair such breaks. One of the non-classical HR mechanism is BIR, which contributes to repair of DSB when only one end shares homology with a donor or one-ended DSBs arise for example at stalled, collapsed or broken replication forks[29–31]. We thus immunostained cells with RAD52, which promotes pathways like alternative-NHEJ and BIR, and observed an increase in RAD52 foci in KMT2C silenced cells along with partial colocalization of RAD52 and 53BP1 foci (Figures 5A, 5B and S4A). Although RAD52 is implicated in a secondary role in presence of RAD51, when HR pathway is compromised as in KMT2C deficient condition, it takes up a key role in repair of collapsed DNA replication forks[32]. We suspect that KMT2C silencing induces DNA breaks in genomic DNA which attempt to get repaired utilizing alternate pathways like BIR.

**Figure 5.**
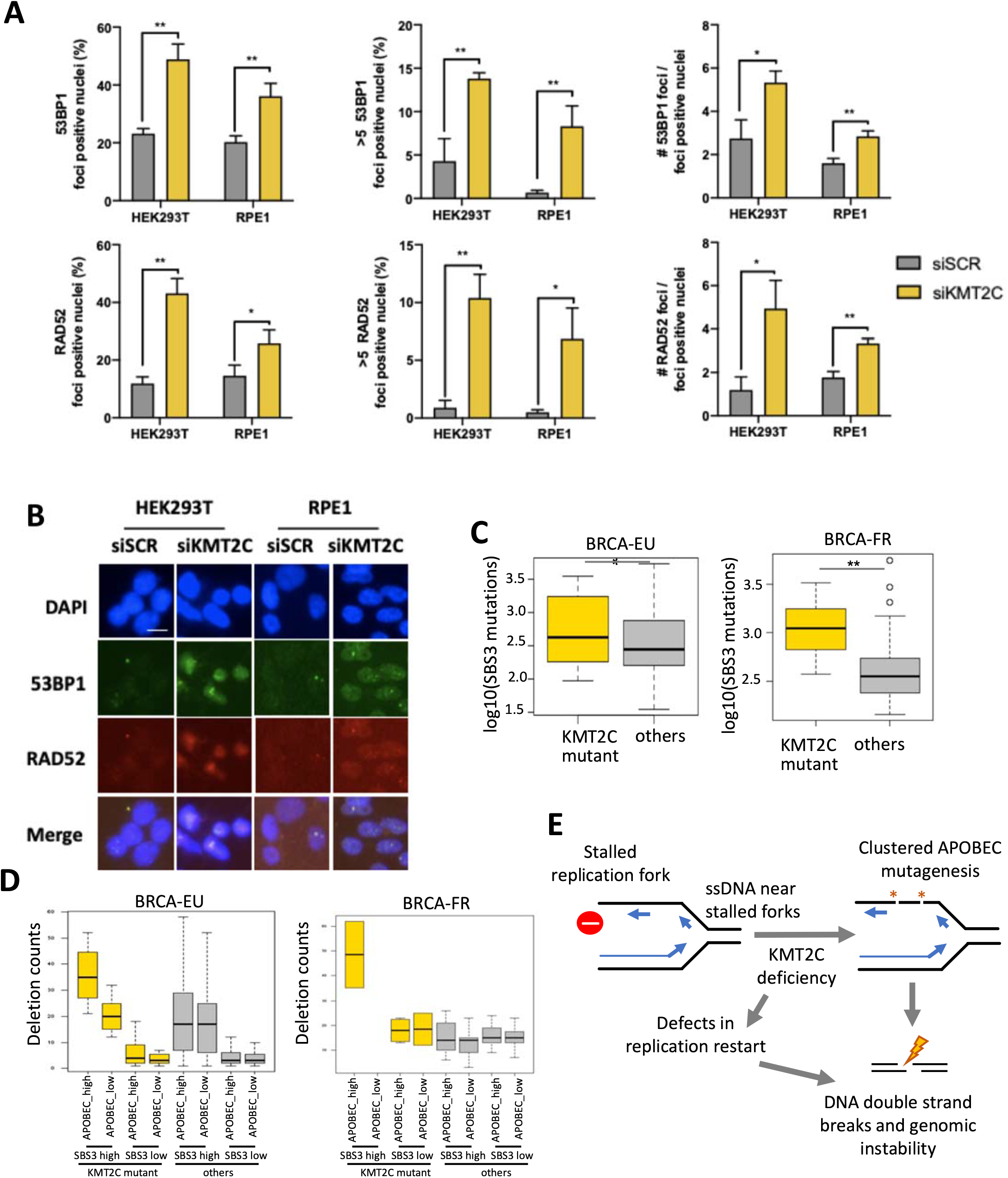
KMT2C mutant tumors have elevated genomic rearrangements and signatures of HR defects. (A-B) Immunofluorescence analysis and representative images show 53BP1 and RAD52 foci upon knockdown of indicated genes in HEK293T and RPE1 cells with error bars representing SEM. P-value was calculated using t-test; *p-value < 0.05, **p-value <0.01. Scale bars indicate 20 μm. DAPI was used as a nuclear counterstain. (C) Barplots showing the burden of somatic mutations attributed to SBS3 in KMT2C mutant and other samples in the breast cancer cohorts BRCA-EU and BRCA-FR. Plots for other cohorts are shown in Figure S4B. P-value was calculated by Mann–Whitney U-test, *p-value < 0.05, **p-value < 0.01. (D) Barplot showing the frequencies of small deletions (<10kb) in tumor samples grouped according to KMT2C mutation status, and the burden of somatic mutations attributed to APOBEC and SBS3 mutational signatures in the breast cancer (BRCA-EU and BRCA-FR) cohorts. The samples with above mean burden of somatic mutations attributed to SBS3 were considered SBS3_high, and otherwise SBS3_low. Likewise, the samples with above mean burden of somatic mutations attributed to SBS2 or SBS13 were considered APOBEC_high, and the samples with below mean burden of SBS2 and SBS13 were designated as APOBEC_low. P-value was calculated using oneway ANOVA. p-value: BRCA-EU: 2.2e-6; BRCA-FR: 3.5e-02. (E) Schematic representation of a model that defects in replication restart and APOBEC mutagenesis promote DNA double strand breaks and genomic instability in KMT2C deficient cells.

Using a complementary approach, we analyzed tumor genomic data from the ICGC cancer cohorts to compare patterns of genomic rearrangements in KMT2C mutant tumors. We found that small deletions (<10kb) were consistently more frequent in KMT2C mutant tumors compared to other tumors in all the breast cancer cohorts analyzed (Figure S4B). We observed consistent results for other cancer cohorts as well (Figure S4B). Since length of homology in structural variation (SV) junction regions was not always available to infer the DSB repair mechanisms for the SVs, we used the single base substitution signature SBS3 as a proxy for HR deficiency-associated genomic instability. In breast cancer cohorts, KMT2C mutant tumors had significantly higher burden of SBS3 (Figure 5C; BRCA-EU: p-value=2.8e-3; BRCA-FR: p-value= 1.2e-3; Mann-Whitney U-test). We observed similar results for other cancers as well (Figure S4C). Taken together, KMT2C deficiency contributes to genomic instability and HR defects in human cancers.

### APOBEC mutagenesis promotes genomic instability in KMT2C deficient tumors

To determine whether APOBEC mutagenesis in KMT2C deficient tumors promotes further genomic instability and favors specific classes of genomic rearrangements, we jointly examined KMT2C mutation status, APOBEC (SBS2 and SBS13) and HR deficiency (SBS3) signatures, as well as the frequency of different classes of genomic rearrangements for ICGC cohorts with available data (Methods). We divided KMT2C-mutant and other tumors into groups that have high or low weight of APOBEC/SBS3 mutational signatures. Small deletion frequencies varied across the groups (Figure 5D; BRCA-EU, p-value=2.2e-6; BRCA-FR, p-value=3.5e-2); both KMT2C mutations and APOBEC activity were associated with increased frequency of small deletions, and the samples with high APOBEC signature and KMT2C mutations had higher burden of deletions than any other category. Extending the analyses to other classes of genomic rearrangements in these cohorts, we observed similar results for small (<10kb) inversions (Figure S4D).

## DISCUSSION

APOBEC mutagenesis is very common in many cancer types. It appears to occur in sporadic, burst-like manner in somatic cells, as observed in lineage-tracing and biochemical experiments[16,33], and often arise late during cancer progression[34]. But its triggers and impact during carcinogenesis are poorly understood. We propose a model that KMT2C deficiency is one of the major factors leads to replication stress and APOBEC activity, which independently and synergistically contribute to mutagenesis and genomic instability in some cancers (Figure 5E).

KMT2C methyltransferase activity imparts H3K4me1 and H3K4me3 enabling MRE11 recruitment at stalled replication forks[8,9,35,36]. KMT2C deficiency slows down replication and delays replication fork restart by compromising TP53 dependent MRE11 recruitment to distressed replication forks. Replication stress in KMT2C deficient condition likely promotes APOBEC3B expression potentially via AP-1 and NF-κB signaling leading to elevated level of deaminase activity. Alongside, KMT2C deficiency also causes delays in replication restart at stalled forks, where ssDNA can act as substrate for APOBEC mutagenesis. Pan-cancer analysis on somatic mutations at TCW trinucleotide contexts and also using COSMIC mutational signatures show that KMT2C mutant tumors have significant excess of APOBEC mutational signatures. We observed excess of mutations at both the RTCA and YTCA contexts, suggesting that perhaps both APOBEC3A and APOBEC3B have elevated activity in KMT2C mutant tumors, but further work needs to be done to firmly establish that.

KMT2C deficiency promotes HR deficiency and genomic instability, and we also observed an increase in 53BP1 and RAD52 foci in KMT2C deficient cells. Increasing evidence suggests that RAD52 gets recruited to stalled forks in distinct HR sub-pathways like BIR, SSA[10,37,38]. Moreover, in yeast 53BP1 also favors the BIR pathway for completion of DNA synthesis[39]. Indeed, we observed partial overlap of 53BP1 and RAD52 foci as well, which suggests that RAD52 becomes essential for maintaining genomic stability and viability of human cells with compromised HR, as in KMT2C deficient conditions. Furthermore, KMT2C insufficiency might promote utilization of non-canonical HR pathways at arrested forks, contributing to the mutational signature SBS3 in human cancers. In KMT2C deficient cells, increased APOBEC activity likely confers further genomic instability by acting on long stretches of ssDNA generated on accumulated stalled forks in KMT2C deficient cells, leading to both point mutations and DNA double strand breaks. Abasic sites generated by APOBEC deaminase activity can also further slow replication fork progression[30], which in turn may facilitate additional mutagenesis, complex processivity patterns, and double strand breaks leading to rearrangements. These processes can promote progressive genomic instability in tumor genomes. While HR deficiency can be targeted by PARP inhibitors[8], APOBEC mutational signatures are considered as potential predictive markers for immunotherapy response[36]. KMT2C is one of the most frequently mutated cancer gene, and occur in older patient populations, often without effective targeted therapies. In such patients, HR defects and APOBEC mutational signatures may help explore opportunities for potentially effective, combinatorial treatment strategies.

## METHODS

### Cancer genomic datasets

We analyzed genomic data including KMT2C mutation status in 7,574 samples from 17 major cancer types from the TCGA [11] and 2,089 samples from 13 cancer types from ICGC [12] cohorts. Different classes of somatic mutations in KMT2C locus were annotated using SnpEff v3.3c[40]. The maftools package[41] and UCSC Genome Browser [42]were used for visualization of mutation spectrum and mutation hotspots in the gene locus. The tumor samples that had at least one non-silent (missense, nonsense, frameshift) mutation that potentially alters the KMT2C gene product were designated as KMT2C-mutant. Several cohorts also reported KMT2C copy number alterations and fusion events, but those appeared to be less frequent compared to point mutations. We also obtained the catalog of somatic structural variations from these ICGC cancer cohorts as well as breast cancer cohort (BRCA-onco) derived from Nones et al.[43].

### Analysis of point mutation and structural variations

We first analyzed the somatic mutations at TCW trinucleotide context (W = A or T; including TCA to TTA or TGA, TCT to TTT or TGT), which are likely due to APOBEC mutagenesis [13]. Since APOBEC3A favored YTCA, while APOBEC3B favored RTCA (Y = pyrimidine, R = purine) contexts [15], we further estimated the prevalence of somatic mutations at these contexts. We also used deconstructSigs [44] with default parameters to determine the proportion of COSMIC v2 mutational signatures in the tumor genomes. For the cancer cohort level analysis, the samples from each of the TCGA or ICGC cohorts were grouped as KMT2C mutant and others, and then mutational patterns were compared between the two groups using an approach similar to that implemented in the mutSigTools[20]. For the pan-cancer analysis, all samples from the TCGA or ICGC cohorts were pooled and grouped based on the KMT2C status as above. For the cohorts with whole genome sequencing data, we analyzed genomic rearrangements as deletions, duplications and inversions, and further grouped the intra-chromosomal rearrangements within 10kb. We further used the burden of somatic mutations attributed to mutational signature SBS3 [45] as a proxy for homologous recombination (HR) mediated DNA repair defects.

### Context-guided analysis of mutational signatures

We annotated the genomic regions based on (i) genomic features - whole genes, exons, repeats, and telomeres, (ii) chromatin features - strong euchromatin, weak euchromatin, intermediate chromatin, weak heterochromatin, strong heterochromatin, (iii) nuclear localization - inter-lamina regions at the nuclear interior, and lamina-associated regions at the nuclear periphery, and also replication context – early and late replication, and compared mutations enrichment of different mutational signatures between KMT2C mutant and other tumors at each context using a published approach [20]. We retrieved ATAC-seq [46]and somatic mutation data from the same TCGA cancer cohorts, and analyzed the cohorts with at least 10 samples. We also obtained data on sites of histone modifications in KMT2C deficient bladder HTB9 (H3K4me1 and H3K27ac) [8]and breast MCF7 (H3K27ac and H3K9ac) [5]cell lines.

We analyzed somatic mutations from BRCA-EU cohort in the context of long-range interactions inferred from HiC data for T47D breast cancer cell line using a published protocol[24]. Libraries were generated using HindIII restriction enzymes, and Illumina HiSeq paired-end reads were aligned using BWA to the human reference genome hg19, duplicate pairs were counted only once, and processed intra- and inter-chromosomal contact matrices were presented at 100kb resolution, as detailed elsewhere (GEO: GSM1294039). Normalized contact score ≥3 between pairs of genomic regions was considered for analyzing patterns of long-range interactions.

### Replication timing analysis

Replication timing data for human cell lines was obtained from the ENCODE project (wgEncodeEH002247). The wavelet-smoothed replication time signal data were divided into deciles (D1, D2,… D10) according to their decreasing ordered values, and equal signals were assigned to each decile. Genomic segments were annotated according to their replication timing decile, such that the earliest and latest replicating regions were mapped to first and last deciles, respectively. Somatic mutations in the breast cancer samples were analyzed in the context of replication timing data of related cell lines (BRCA-EU mutation data was analyzed with replication timing data from MCF7 breast cancer cell line). Somatic mutations within genomic regions corresponding to different replication decile were identified and mutations enrichment of corresponding COSMIC mutational signatures were calculated using published approaches[17]. Several mutagenic processes (including APOBEC mutagenesis) results in clusters of mutations on the same DNA strand, which is quantified by processivity. Processive groups were determined as maximal stretches of adjacent mutations containing the same reference alleles and were generated by the same signature in each sample. The mutational signature of each substitution was assigned to the highest a posteriori probability, which was identified using the quadratic programming (QP) approach[47]. Kataegis events were filtered to decrease bias signals, which did not alter the key conclusions.

### Regulation of APOBEC expression

We obtained data on regulatory regions and transcription factors (TFs) binding sites located around APOBEC3B locus in the U2OS cell line from a published report [21]. Primarily TFs were from NF-κB gene family (NFKB1, NFKB2, RELA, RELB) and from the AP-1 gene family (FOS, FOSL1, FOSL2, JUN, JUNB, JUND) were shown to co-regulate APOBEC3B expression. We also obtained the processed ATAC-seq peak calls [46] and APOBEC3B expression data for the TCGA cancer cohorts [11] with sufficient number of KMT2C proficient and deficient samples (TCGA-BRCA and TCGA-COAD).

### Cell culture and knockdown of KMT2C expression

HEK293T, RPE1 and MDA-MB-231 cells were cultured in Dulbecco’s modified eagle’s medium (D6429, Sigma-Aldrich), and SkBr3 in Dulbecco’s modified eagle medium/Nutrient mixture F-12 (11320033, GIBCO) supplemented with 10% fetal bovine serum (97068085, VWR) and 1% Penicillin-Streptomycin solution (97063708, VWR), at 37°C in a humidified incubator with 5% CO_2_, and harvested using trypsin-EDTA solution (25200056, GIBCO) or rubber-tipped cell scraper. SMARTpool siRNAs were used for knockdown of KMT2C (siKMT2C) (L-007039, Horizon Discovery) and Scrambled (siSCR) was used as a negative control (D-001810, Horizon Discovery). For protein knockdown, cells were transfected with the indicated siRNAs at a final concentration of 50 nM using Lipofectamine (13778075, Thermo Fisher Scientific) following standard protocols for cell lines. After incubating the cells for 48 h, 5 mM HU (400046, Sigma-Aldrich) was added where indicated and the cells were incubated for an additional 1 h.

### Measurement of gene expression by real-time quantitative PCR (RT-qPCR)

Total RNA from cells was isolated using Trizol (15596026, Thermo Fisher Scientific) as per manufacturer’s instructions. 1μ of total RNA was reverse transcribed using High-Capacity cDNA Reverse Transcription Kit (4368814, Thermo Fisher Scientific). One-tenth of this cDNA was used per real-time PCR assay and each sample was assayed in triplicates. Power SYBR green master mix (4367659, Thermo Fisher Scientific) was used for RT-qPCR and samples were quantified on QuantStudio™ 12K Flex Real-Time PCR System (Thermo Fisher Scientific). In brief, CT values using gene specific primers were normalized to the CT value of actin. Fold change in gene expression across samples was calculated by the formula 2^-ΔΔCT^. Gene expression values from control experiments were set to unity and data from all treatments were calculated as fold change over 1. All assays were done using RNA from three independent experiments unless otherwise stated. Primers used were - KMT2C (F): TTA CAC ACA GTG CGC TCC TT, KMT2C (R): AGG GTC TGC ACA TGC TAC AA; MRE11 (F): CAG TGT TTA GTA TTC ATG GCA ATC ATG, MRE11 (R): AAT GTC CAA GGC ACA AAG TGC; APOBEC3B (F): GAC CCT TTG GTC CTT CGA C, APOBEC3B (R): GCA CAG CCC CAG GAG AAG, ACTIN (F): GAG CAC AGA GCC TCG CCT TT, ACTIN (R): TCA TCA TCC ATG GTG AGC TGG. The experiment was performed at least thrice and data analyzed using two-tailed t-test. Values in the plot are means ± SEM.

### DNA fiber assay

DNA fiber assay on Scrambled siRNA and KMT2C siRNA treated HEK293T cells was performed. Experiments performed with or without hydroxyurea (HU) treatment under conditions indicated in the schematic. Briefly, asynchronous HEK293T cells were labelled with 25 μM CldU for 20 min, washed with PBS, treated or not with 2 mM HU for 2 h, washed again in warm PBS and exposed to 250 μM IdU for 20 min before collection. Cells were then lysed and DNA fibers stretched onto glass slides. The fibers were denatured with 2.5 M HCl for 1 h, washed with PBS and blocked with 2% BSA in phosphate buffered saline Tween-20 for 30 min. The newly replicated CldU and IdU tracts were incubated with anti-BrdU antibodies recognizing CldU and IdU respectively (ab6326, Abcam; 347580, BD Biosciences) for 2 h at room temperature in a humidified chamber. Fibers were then washed with PBS and incubated with secondary antibody against CldU (A-11007, Thermo Fisher Scientific) and IdU (A-11029, Thermo Fisher Scientific) for 1 h at room temperature in a humidified chamber. After washing in PBS, the slides were air-dried and mounted using VECTASHIELD Mounting Medium (H-1200, Vector Laboratories). Images were taken at 40x with oil immersion objective using a Nikon Eclipse TE2000-U microscope (Nikon). Images of the same group were captured with identical exposure time using NIS-Elements software and analyzed using ImageJ software. The length of minimum one hundred fifty tracts from each condition were measured. IdU/CldU values were analyzed using Mann-Whitney U-test with median and interquartile range shown. Fork stalling/ new origins firing was estimated by analyzing red tracts only for stalled replication forks and green only and green–red–green for newly fired origins, using two-tailed t-test with more than 250 fibers measured for each condition. The experiment was performed at least twice.

### Quantification of 53BP1 and RAD52 double immunostained foci

Cells were washed with PBS following treatments, fixed with 3% paraformaldehyde and permeabilized with 0.5% Triton X-100. Cells were blocked with 1% BSA for 1 h, then incubated sequentially with primary antibodies (53BP1-ab21083, Abcam; RAD52-MA5-31888, Thermo Fisher Scientific) and secondary antibodies (A11008, Thermo Fisher Scientific; A32742, Thermo Fisher Scientific respectively) for 1 h each at 37°C, with three PBS washes in between. Coverslips were mounted onto glass slides with VECTASHIELD Mounting Medium with DAPI (H-1200, Vector Labs). Images were captured at 20x objective using EVOS M500 microscope (Thermo Fisher Scientific). Images of the same group were captured with identical exposure time. Images were processed using ImageJ software, and cells were scored as displaying either diffused or punctuated staining. Cells with punctuated staining were further analyzed for calculation of the number of foci. The experiment was performed at least thrice and data analyzed using twotailed t-test. Values in the plot are means ± SEM.

### Preparation of protein lysate and cytidine deamination assays

Cells were lysed in HNET buffer (25 mM HEPES-NaOH, 150 mM NaCl, 5 mM EDTA, 1% Triton X-100, 1 mM DTT, 1x Protease Inhibitor Cocktail (P2714, Sigma-Aldrich). Protein lysates were quantified by BCA method (PI23227, Thermo Fisher Scientific) and equal amounts were used for subsequent assays. Deaminase assay was performed as described by [48]. Briefly, equal amounts of protein lysate were incubated with 5 pmol probe in 1x deaminase buffer (10 mM Tris.HCl pH 8.0, 50 mM NaCl, 1 mM DTT) at 37°C for 2 h followed by incubation with 0.75 U UDG (EN0361, Thermo Fisher Scientific) in 1x UDG buffer at 37°C for 45 min. The reaction mixture was treated with 0.15 M NaOH at 37°C for 20 min prior to its termination by heating at 95°C for 3 min with equal volume of 2x RNA loading dye and prompt chilling on ice. Samples were resolved by electrophoresis at 150 V for 3 h at room temperature in Novex TBE-Urea Gels (EC6885BOX, Thermo Fisher Scientific) in 1x TBE buffer and cleaved and uncleaved products were imaged on ChemiDoc (Bio-Rad) using Alexa Fluor 488 module. Percent substrate cleaved was determined by quantification of the intensity of the substrate and cytidine deamination cleavage product using ImageJ (NIH). Fluorescently labeled DNA probes used for the assay were - Test probe: 5Alex488N/AT AAT AAT AAT AAT AAT AAT AAT ATC CAT AAT AAT AAT AAT AAT AAT A; Positive probe: 5Alex488N/AT AAT AAT AAT AAT AAT AAT AAT ATU UAT AAT AAT AAT AAT AAT AAT A. The experiment was performed at least thrice, and data analyzed using two-tailed t-test. Values in the plot are means ± SEM.

### Statistical analyses

Statistical analyses of genomic data were performed using R version 3.6.1 and Prism 8 (GraphPad). Asterisks indicate statistical significance wherein **p < 0.01; *p < 0.05; ns, non-significant. For DNA fiber assay we used more stringent test with additional cutoffs with ****p < 0.0001, ***p < 0.001. Statistical tests and corresponding p-values are listed for respective analyses.

## ACKNOWLEDGEMENTS

The authors acknowledge financial support from R01GM129066, P01CA250957, Robert Wood Johnson Foundation (SD). The authors thank Yizhou He for the RPE1 cell line, Michael L. Gatza for HEK293T, Dr. Bing Xia for MDA-MB-231 cell line, Dr. Hatem Sabaawy for SkBr3 cell line, and Vinod Singh for input regarding mutSigTools. We thank Dr. Bing Xia’s lab for assistance and helping with chemicals for DNA fiber assay.

## AUTHOR CONTRIBUTION

SD conceived the project. XH, AB, and SD designed the experiments. AB performed the experiments. XH, AB and SD analyzed the data. XH, AB, and SD wrote the paper.

### Declaration of interests

The authors declare no competing interests.

## SUPPLEMENTARY INFORMATION

Supplementary Figures S1-S4.

## Notes

### Competing Interest Statement

The authors have declared no competing interest.

