## Supplementary Figure for "KMT2C deficiency promotes APOBEC mutagenesis and genomic instability in multiple cancers"

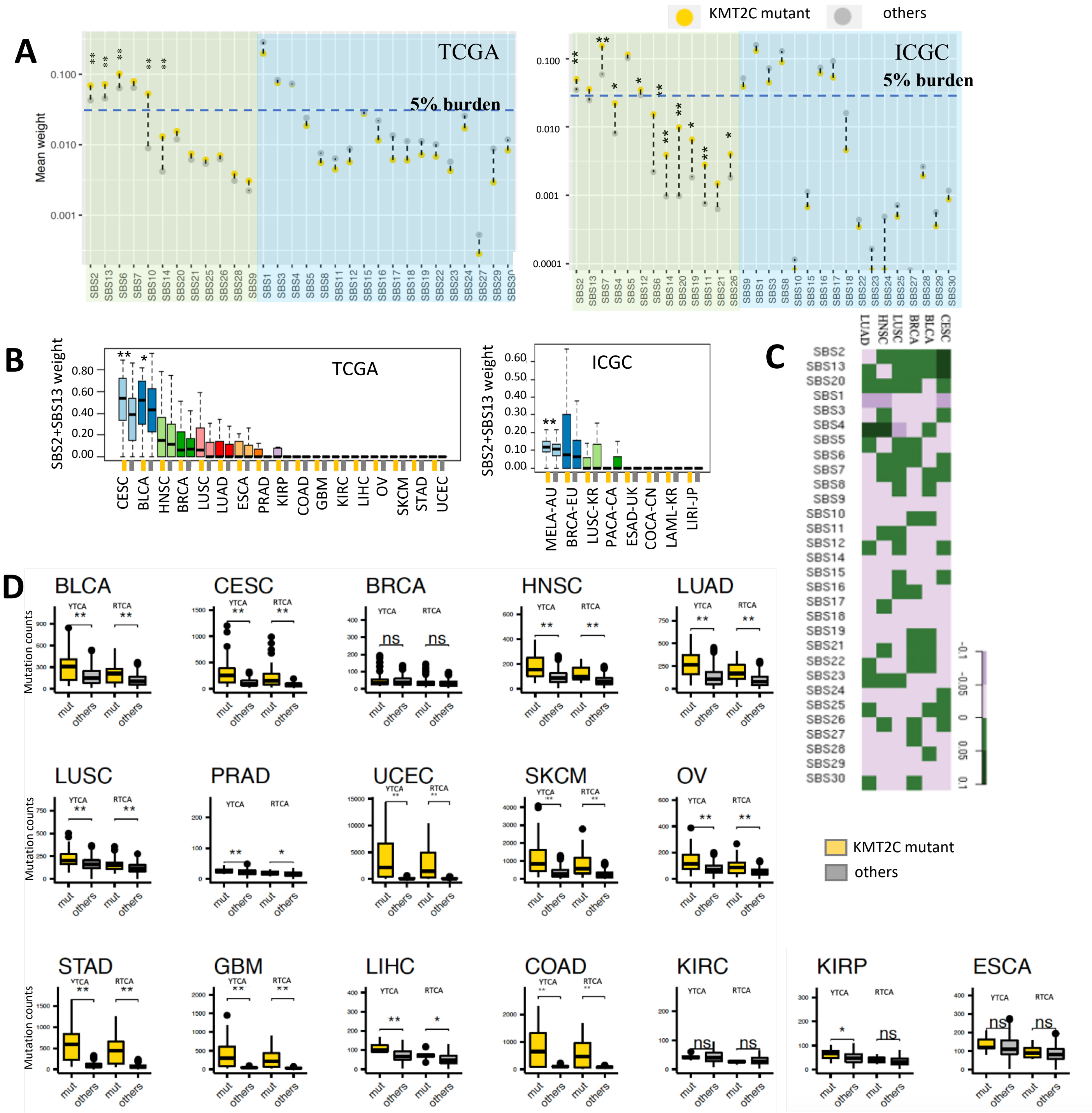


**Figure S1 (related to Figure 1). Burden of APOBEC mutational signatures across TCGA and ICGC cohorts. A**) Distribution of the weights of the COSMIC v2 mutational signatures (SBS1-30) in KMT2C mutant and other samples in a pan-cancer analysis using the TCGA and ICGC datasets. P-value was calculated by Mann–Whitney U-test, *p-value < 0.05, **p-value < 0.01. **B)** Distribution of the weights of the APOBEC mutational signatures (SBS2 and SBS13) in the KMT2C mutant and other samples in the TCGA and ICGC cancer cohorts. P-value was calculated by Mann–Whitney U-test, *p-value < 0.05, **p-value < 0.01. **C)** The heatmap showing relative preference for the 30 COSMIC signatures in the KMT2C mutant tumors relative to the other samples in 6 cancer cohorts. Relative preference for i-th mutational signature (i=1, 2, .. 30) was computed as ($M_i^{KMT2C\ mut}$ - $M_i^{others}$), $M_i$: mean weight of i-th mutational signature. **D)** The count of somatic mutations at RTCA (including ATCA and GTCA) and YTCA (including CTCA and TTCA) contexts in the KMT2C mutant(yellow) and other (grey) samples in TCGA cancer cohorts. P-value was calculated by Mann–Whitney U-test, *p-value < 0.05, **p-value < 0.01.

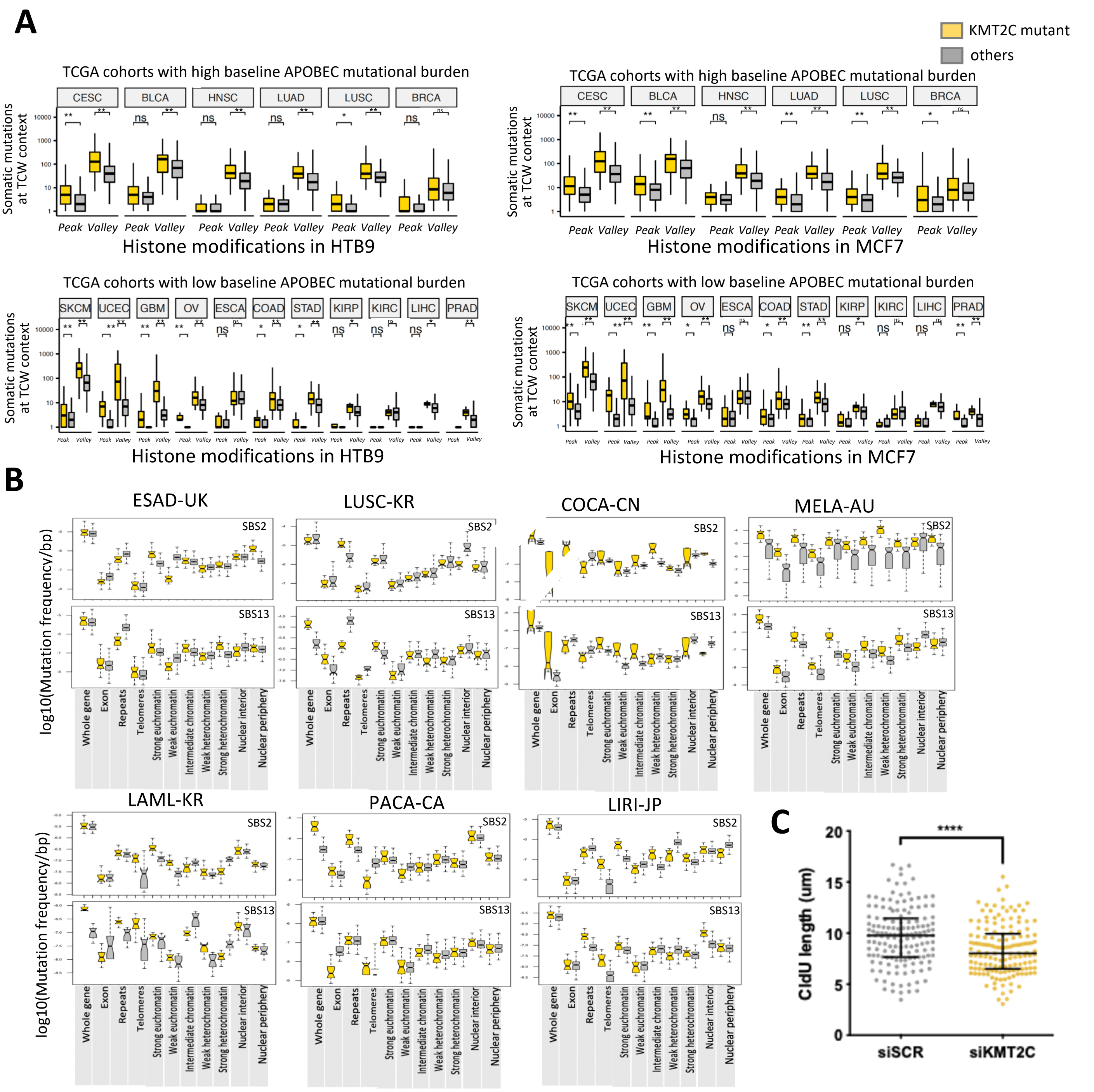


**Figure S2 (related to Figure 2). Context-dependent preferences for APOBEC mutational signatures in KMT2C mutant tumors. A)** Boxplots showing somatic mutation frequencies at TCW trinucleotide contexts, grouped according to their location within regions of peak or valley regions in KMT2C mutant and other samples in respective cancer cohorts. Open (peak) or closed (valley) chromatin regions defined as the sites of histone modifications (H3K4me1 and H3K27ac for HTB9 cell line and H3K4me3, H3K27ac and H3K9ac for MCF7 cell line). P-value was calculated by Mann–Whitney U-test, *p-value < 0.05, **p-value < 0.01. **B)** Enrichment of APOBEC mutation signatures SBS2 and SBS13 at different genomic, epigenomic, and nuclear localization contexts in KMT2C mutant(yellow) and other samples (grey) in ICGC cancers. **C)** DNA fiber assay on control (siSCR) and KMT2C knockdown (siKMT2C) HEK293T cells indicating CldU lengths with line and bars representing the median and interquartile range. Asterisks indicate statistical significance.

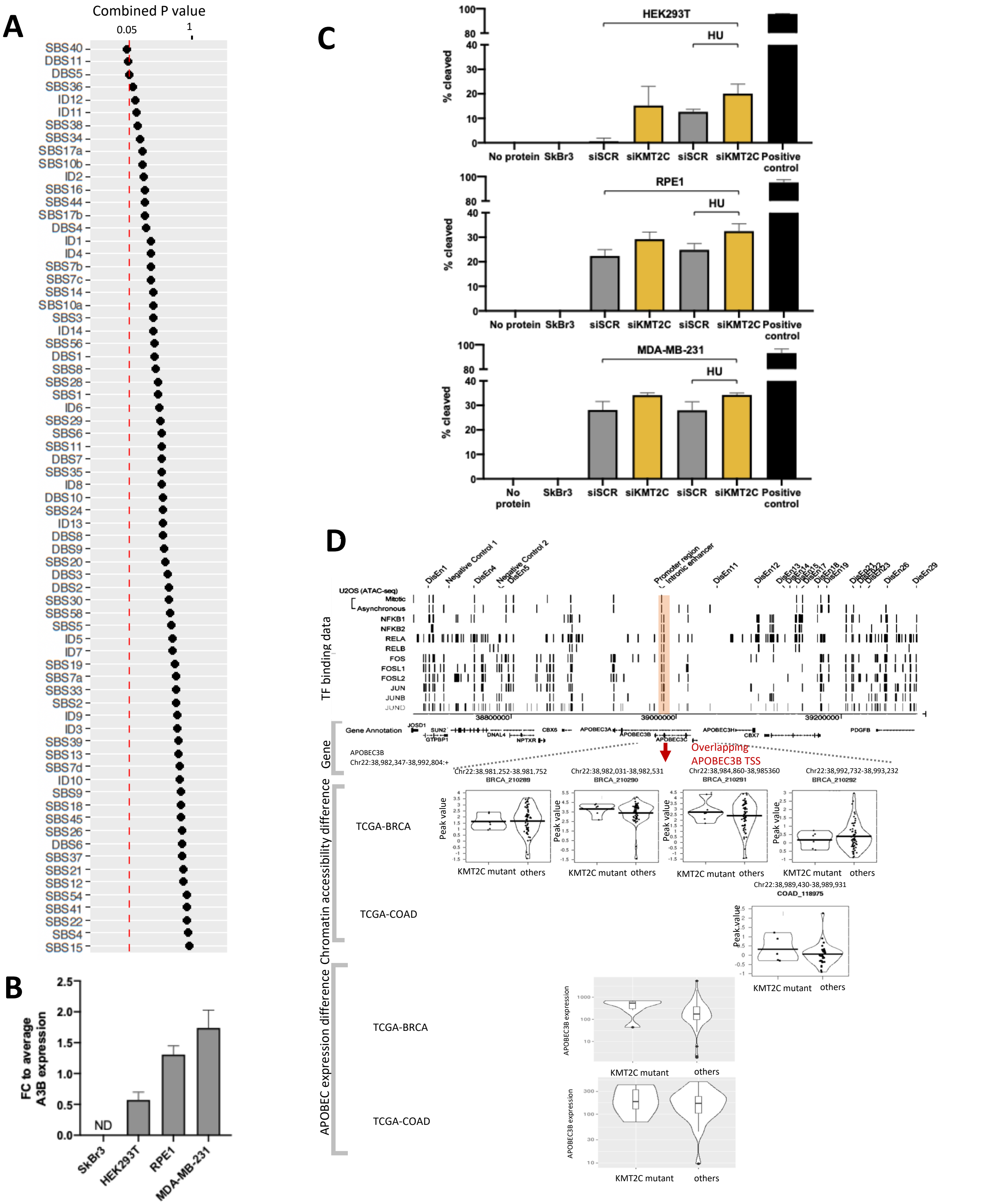


**Figure S3 (related to Figure 3). KMT2C loss induces APOBEC expression and activity. A)** Combined p-values showing the significance of regression coefficients of the burden of single (SBS) and double base substitution (DBS) signatures (COSMIC v3.1) against KMT2C and MRE11 expression in the PCAWG samples. **B)** Fold change (FC) in mRNA expression of APOBEC3B (A3B) in the indicated cell lines as determined by quantitative PCR with error bars representing SEM. ND indicates non-detected. **C)** Percentage of cleaved probe in deaminase assay using lysates from indicated cell lines for shown conditions with no protein and positive probe only were used as negative and positive controls, respectively with error bars representing SEM. P-value was calculated using t-test wherein *p-value < 0.05, **p-value < 0.01. **D)** (Top) Known transcription factor binding sites, (middle) cancer type-specific ATAC-seq peak regions in the TCGA cancer cohorts, and (bottom) APOBEC3B gene expression in the TCGA cancer cohorts with matched RNA-seq, ATAC-seq, and KMT2C mutation data are shown. KMT2C mutant samples have more open chromatin around APOBEC3B promoter region in the TCGA breast (BRCA) and colon (COAD) cancer cohorts, and this difference is most prominent in the ATAC-seq peaks around APOBEC3B TSS. These peaks are proximal to the binding sites of the NF-κB and AP-1 family, which are reported to regulate APOBEC3B expression (Lin et al., 2020). KMT2C mutant samples with more open chromatin around APOBEC3B promoter region had relative increase in APOBEC3B expression relative to the KMT2C proficient tumors in both the BRCA and COAD cohorts, but sample size and p-values were modest (p-value >0.05; Wilcox test).

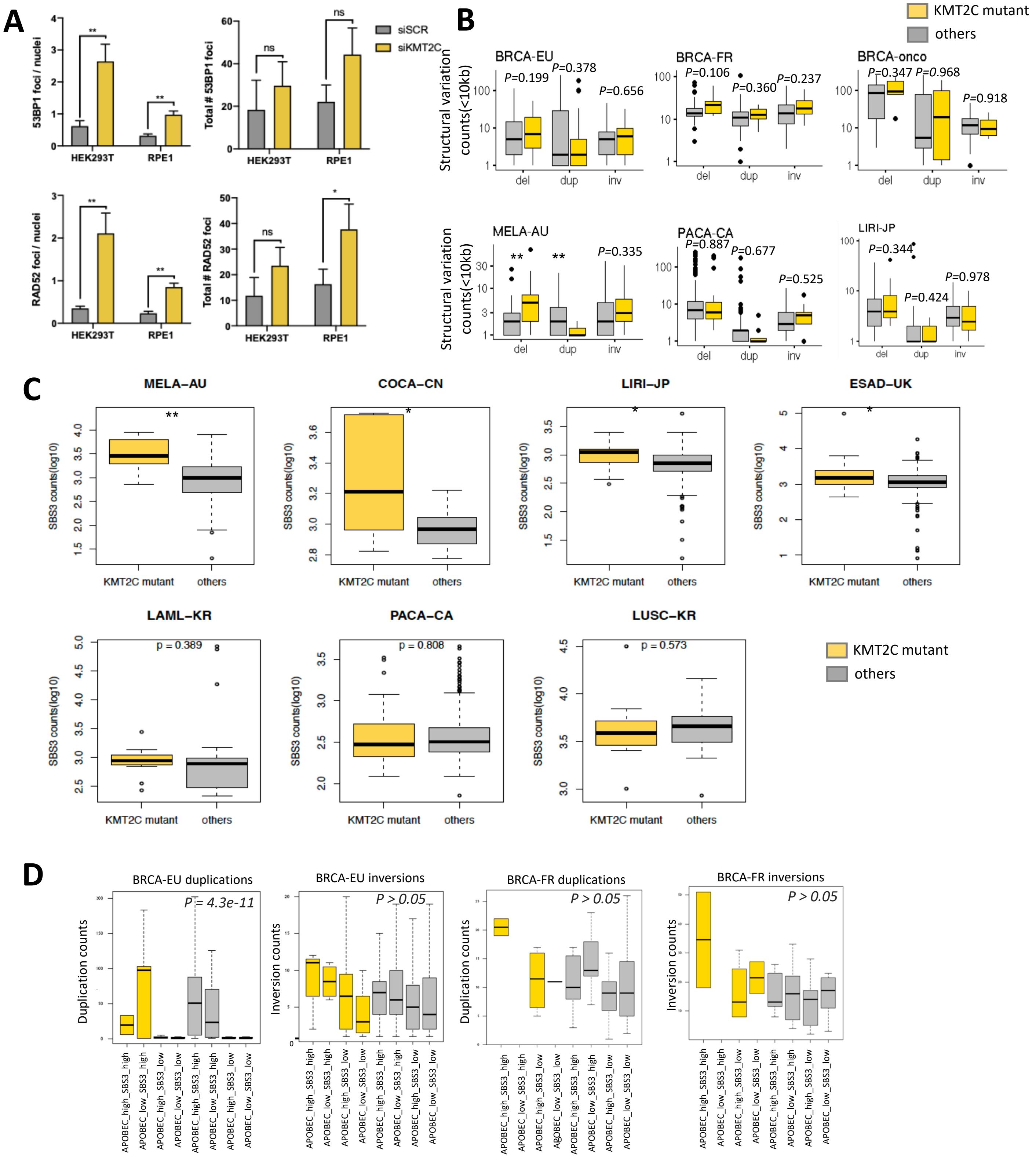


**Figure S4 (related to Figure 5). KMT2C loss promotes genomic instability in cancer genomes**. A) Quantification of 53BP1 and RAD52 foci upon knockdown of indicated genes in HEK293T and RPE1 cells with error bars representing SEM. P-value was calculated using t-test wherein *p-value < 0.05, **p-value < 0.01. B) Boxplot showing the frequency of small (<10kb) structural variations (deletions, inversions and duplications) in KMT2C mutant and other samples in ICGC and BRCA-onco (Nones et al., 2019) cohorts. P-value was calculated by Mann-Whitney U-test. *p-value < 0.05, **p-value < 0.01. Del: deletion, dup: duplication, inv: inversion. C) Barplots showing the number of somatic mutations attributed to SBS3 in KMT2C mutant and other samples in ICGC cancer cohorts. P-value was calculated by Mann–Whitney U-test, *p-value < 0.05, **p-value < 0.01. D) Barplot showing the frequency of small duplications and inversions (<10kb) in samples grouped according to the KMT2C mutation status, and the burden of SBS3 and APOBEC mutation signatures in breast cancer cohorts, as described in Figure 5. BRCA-onco cohort did not have sufficient data for an equivalent analysis. The p-value was calculated using one-way ANOVA.
